# Nanopore Sequencing in Microgravity

**DOI:** 10.1101/032342

**Authors:** Alexa B.R. McIntyre, Lindsay Rizzardi, Angela M Yu, Gail L. Rosen, Noah Alexander, Douglas J. Botkin, Kristen K. John, Sarah L. Castro-Wallace, Aaron S. Burton, Andrew Feinberg, Christopher E. Mason

## Abstract

The ability to perform remote, in situ sequencing and diagnostics has been a long-sought goal for point-of-care medicine and portable DNA/RNA measurements. This technological advancement extends to missions beyond Earth as well, both for crew health and astrobiology applications. However, most commercially available sequencing technologies are ill-suited for space flight for a variety of reasons, including excessive volume and mass, and insufficient ruggedization for spaceflight. Portable and lightweight nanopore-based sequencers, which analyze nucleic acids electrochemically, are inherently much better suited to spaceflight, and could potentially be incorporated into future missions with only minimal modification. As a first step toward evaluating the performance of nanopore sequencers in a microgravity environment, we tested the Oxford Nanopore Technologies MinION™ in a parabolic flight simulator to examine the effect of reduced gravity on DNA sequencing. The instrument successfully generated three reads, averaging 2,371 bases. However, the median current was shifted across all reads and the error profiles changed compared with operation of the sequencer on the ground, indicating that distinct computational methods may be needed for such data. We evaluated existing methods and propose two new methods; the first new method is based on a wave-fingerprint method similar to that of the Shazam model for matching periodicity information in music, and the second is based on entropy signal mapping. These tools provide a unique opportunity for nucleic acid sequencing in reduced gravity environments. Finally, we discuss the lessons learned from the parabolic flight as they would apply to performing DNA sequencing with the MinION™ aboard the International Space Station.

## Introduction

Remote molecular diagnostics on Earth and in human space exploration missions will require portable technologies for real-time evaluation of environmental microorganisms and crew health^1,2^. Such technologies could prove critical for rapid responses to medical infections in space, including for determining whether or not antibiotics are needed for an infection, or which set of antibiotics to use. Single-molecule methods can also enable the discovery of modified nucleic acids, a potentially important aspect to ongoing monitoring of crew health^3,4^. This capability could equally benefit the search for life beyond Earth by expanding the range of detectable information polymers beyond the canonical nucleobases of DNA and RNA bases.

The MinION™ sequencer from Oxford Nanopore Technologies (ONT) is a sequencing device approximately the size of a large USB stick (4ʺ×1.5ʺ×1ʺ, with a mass of ~100g) that draws power from and transmits data to a computer through a single USB 3.0 connection. The combination of a nanopore sequencer and lightweight tablet computer offers a heretofore-unseen portability in nucleotide sequencing. Two advantages of nanopore sequencing are a relatively straightforward library preparation protocol and the absence of an amplification step prior to sequencing. DNA molecules are prepared through the ligation of a hairpin adapter to one end of the double stranded molecule, joining the template and complementary strands. This structure permits sequencing of both sides (2D) of the library templates, derived when leading strand, hairpin adapter, and complementary strand pass through the pore in succession. The increase in information stored in such 2D reads produces more accurate base calls than reads of template or complement strands alone.

Although the technology continues to improve, significant challenges remain in the production and interpretation of nanopore sequencing data, due to error rates of ~15% for 2D reads^5^. However, for applications such as microbial identification, the long reads, on the order of several thousands of nucleotides or more, are frequently sufficient to permit classification at the species or genus levels. Considering the portability of MinION™ sequencer and the powerful organism identification capability offered by sequencing, preparations have begun for the MinION™ sequencing platform, a Microsoft Surface Pro 3 tablet for power and data handling, and ground-prepared samples to be used for sequencing aboard the International Space Station in the summer of 2016. Here, we describe a first look at our in-flight concept of operations (sample loading, tablet set-up, etc.) and the performance of the MinION™ sequencer under the intermittent microgravity conditions that occur on a microgravity parabolic flight. Although the present experiments were performed with sequencing libraries that were prepared on the ground, we posit that library preparation could also be performed in space with the liquid handling procedures described in Rizzardi et al. (2015).

Successful detection of nucleotides depends on the ability of the bases in the pore at a given time to disrupt ionic current flow with a specific signal dependent on their identity. By calling events at time points where raw electric current measurements change significantly, event data can be used to infer the sequence of a polynucleotide chain. Thus, each event time point should ideally reflect the entry of a single new nucleotide into the pore. However, the current detection process remains subject to a high degree of noise and a strong dependence on reaction conditions. ONT currently uses a hidden Markov model (HMM) algorithm with a Viterbi decoder algorithm by Metrichor™ to call bases from event data^7^, but traditional alignment software has failed to map most reads produced through this pipeline^8,9^. Thus, nanopore data provides a novel challenge to base-calling and analysis. We discuss the performance of several existing methods for classifying nanopore reads by species using data from the microgravity flight and a control experiment performed on the ground, as well as present two novel methods for analysis using noisy event squiggle data from current measurements.

## Results

### Testing the MinION^TM^ DNA sequencer in microgravity

As a further proof-of-principle for our study regarding liquid handling in microgravity^6^, we demonstrate the possibility of performing an actual genomics-level experiment in space using the MinION^TM^ sequencer. Pre-filled syringes (with plastic pipette tips) containing ready-to-load DNA libraries were prepared for loading the nanopore sequencer during the parabolic flight (see *Methods*). The flow cell was loaded into the MinION^TM^ prior to parabolic flight (Figure 1a).

**Figure 1:**
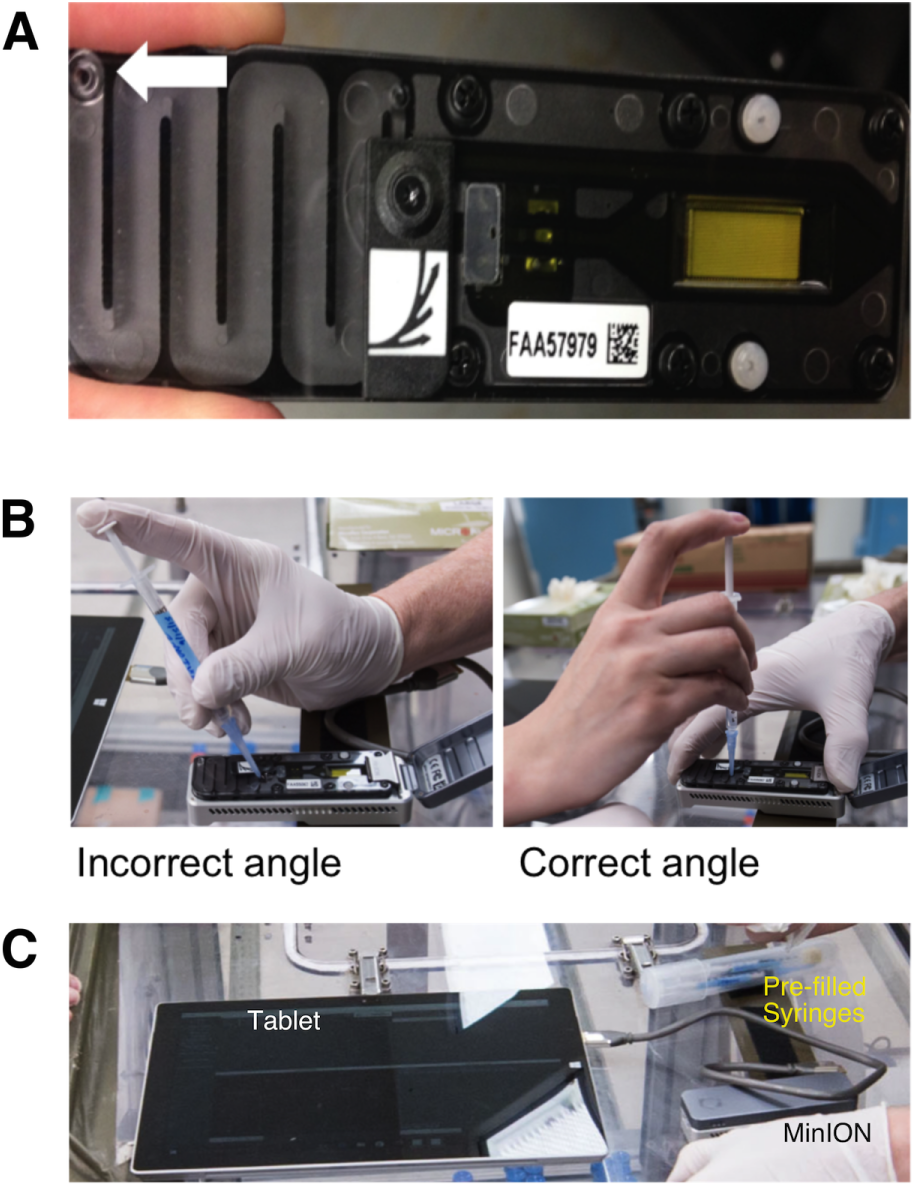
**a)** The MinION™ flow cell, which was loaded into the device prior to the parabolic flight. The white arrow marks vents, which leaked during return transport to the Johnson Space Center. **b)** Loading the library onto the MinION™. Angling the pipette perpendicular to the pore was necessary to avoid introducing air bubbles. **c)** The MinION™ setup on the plane. The flow cell was connected to a tablet running Oxford Nanopore Technologies’ MinKNOW™ sequencing software via a USB 3.0 cable. We noted significant glare off of the tablet screen.

During the microgravity portion of parabolic flight, we performed air bubble removal and loaded the DNA library mixture into the flow cell. An empty syringe was used to remove the initial air bubble from the input pore of the flow cell. We did not observe any bubbles in the flow cell pre-flight. To prevent introduction of an air bubble, the library mixture was pushed to the very end of the tip before engaging with the sample-loading pore. it was critical when removing the air bubble and loading the DNA library that the pipette tip be firmly engaged and held exactly perpendicular to the sample loading pore (Figure 1b). This creates a seal, allowing introduction of the sample without including an air bubble. After library loading, the flow cell was connected via a USB 3.0 cable to a Microsoft Surface Pro 3 tablet running the sequencing software (Figure 1c).

Sequencing was initiated immediately after sample loading and continued for the remainder of the flight (10 parabolas of approximately 1 minute each, including 30 seconds of microgravity) and through transport back to Johnson Space Center. Data were then collected and transferred to the Mason laboratory for analysis. After halting and opening the sequencer, we observed that the flow cell had leaked from a vent (Figure 1a, arrow) likely due to being tilted vertically during transport to the Johnson Space Center. We did not observe any fluid leaks during parabolic flight.

The number of available nanopores for sequencing during the parabolic flights was much lower than during normal terrestrial sequencing. The first group had 16 available pores, while the remaining three groups had zero active pores. There are several possible reasons for this, including that the flow cells experienced multiple transitions from 1.8 g to microgravity during, both prior to sample loading, as well as during the Mux scan for the four groups. In addition, the flow cells themselves are optimally used within 8 weeks of their receipt, while the flow cell used for the parabolic flight was approximately 12 weeks old. Despite these difficulties, the MinION^TM^ successfully sequenced three reads of varying lengths.

### MinION^TM^ data analysis

Two MinION™ runs were prepared using the same sample, the first run over the course of the parabolic flight and the second entirely on the ground. The sample contained approximately equal masses of DNA from three species: Bacteriophage Lambda (cI857*ind* 1 *Sam* 7), *Escherichia coli* (K12 MG1655), and mouse (BALB/C female genomic DNA). Despite the technical issues described above, the library injected during the parabolic flight produced three template strand reads from two MinION™ channels, with the longest produced at least partially in microgravity, although the time stamps of the other two indicate they were generated after parabolas had concluded. The ground control experiment produced 1,737 template strand (1D) reads from 261 channels, as well as 1,012 complement strand reads and 196 2D pass-filter reads, in approximately 3 hours. For direct comparisons between the two experiments, we examined only the template strands (1D) of the ground data unless otherwise noted.

One of the most salient differences between the signal produced entirely on the ground and that from the MinION™ sent on the parabolic flight is the median current levels across all possible sequences of five bases (5 nucleotide kmers). The flight data exhibited increased currents, with a median of 91.77 pA across reads, while the ground data exhibited a median of 74.8 pA (Figure 2a). These data represent mean shifts from the median currents stored in the HMM of 32.1 and 10.8 pA respectively. Both data sets still produced roughly the correct distribution of amperages across different kmers.

**Figure 2:**
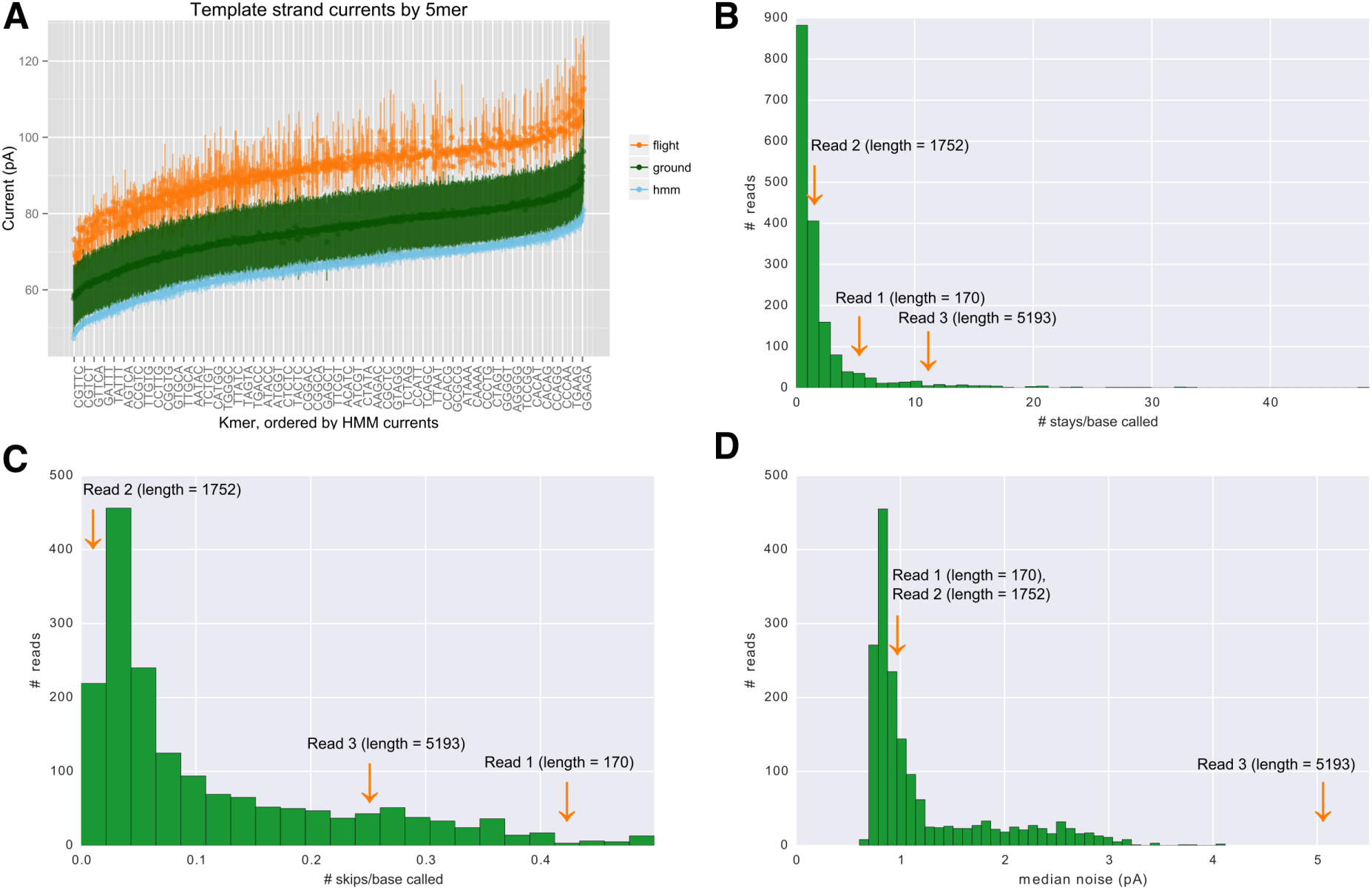
**a)** Current distributions by 5-mer for flight data, ground data, and Metrichor^TM^’s Hidden Markov Model. Standard deviations of the mean were calculated across reads for the flight and ground data, while the stored standard deviation was taken for the HMM 5-mers. The two measures are not directly comparable. **b-d)** Histograms of the number of stays per base, number of skips per base called, and noise for ground data reads. The orange arrows mark the approximate positions of the flight data reads. Median noise is calculated by MinKNOW™ for each read.

Various measures of read quality were also assessed. The ONT event calling software MinKNOW™ produced signals of 977, 3710, and 63,362 events long from the flight run, but base calling generated reads from these signals of only 170 bases, 1,752 bases, and 5,193 bases respectively. MinKNOW™ defines “events” at current changes large enough to suggest that a new base has entered the pore. The base calling algorithm by Metrichor™ then determines how the sequence of these events should translate to the sequence of nucleotide bases. The ratio of the number of events to the number of bases reveals a high proportion of “stays” in the flight data, particularly in the final read. A stay is when an event determined from the amperage signal does not correspond to new bases in the predicted sequence. The average number of stays per base called was higher in the flight data reads with a mean of 5.97, as compared to 2.11 for the ground data (Figure 2b). “Skips” in the signal, bases predicted that do not correspond to events, occurred at a much lower rate than stays in both data sets, with a mean rate of 0.24 skips/base called for the flight data, and 0.11 skips/base called for the ground data (Figure 2c).

The failure to translate over 90% of events to bases in the longest read suggests a high degree of noise. Indeed, the median current noise level as measured by MinKNOW™ in for the longest flight data read and the only read produced during the parabolas (5.04 pA) was higher than in any of the ground data reads, and the other two flight data reads demonstrated more moderate levels of 0.94 and 0. 91 pA respectively (Figure 2d). For comparison, ground template strands featured a median noise level of 0.92 pA across reads.

We ran the Basic Local Alignment Search Tool (BLAST) using blastn settings on the base-called reads to evaluate species detection from the mixed sample^10^. The shortest of the three reads did not map to any species in the sample, while the longest aligned to multiple mammalian species including mouse and human but with only 8% query coverage for the top mouse hit. The medium-length read mapped moderately well to *Escherichia coli*, with 67% identity and 92% query coverage. Blast results for the template strands were typically poor, with average identity of 78% but only 37% query coverage (Figure 3a). Almost a third of reads did not map to any of the three sample species at all. However, running BLAST on the 2D ground data reads returned 55 reads as *Lambda phage*, 72 as *Escherichia coli*, 51 as mouse, 17 as exclusively other species, and 1 as none (Figure 3b). The mean identity for 2D hits was 84%, with mean query coverage of 73%.

**Figure 3:**
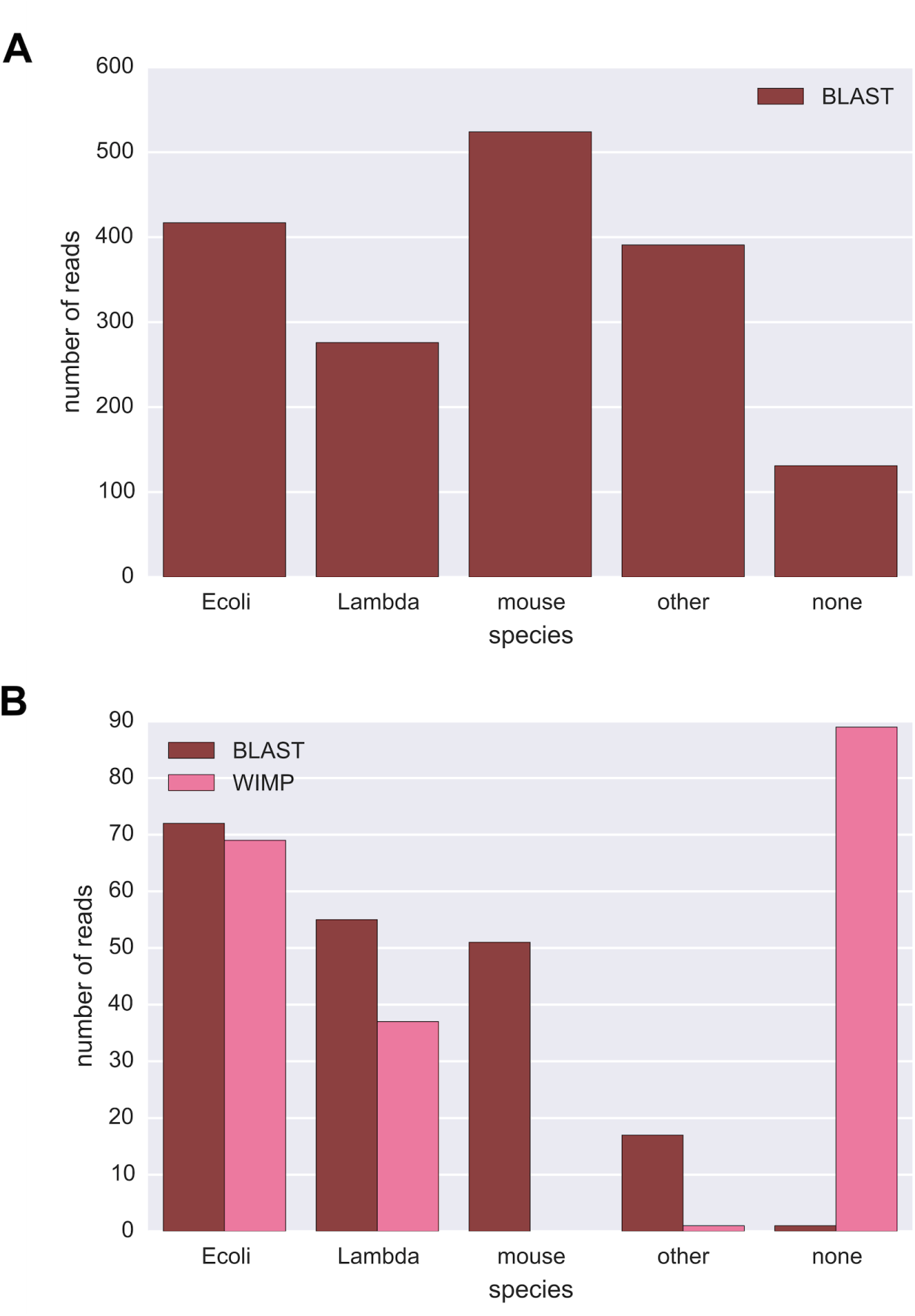
**a)** Blastn results for template strands from the ground control experiment. BLAST hits were preferentially counted towards the sample species even if others scored higher. In the case of ambiguity between multiple sample species, only the highest scoring was considered a hit. **b)** A comparison of BLAST and Oxford Nanopore Technologies’ WIMP using the 2D reads from the ground experiment. We ran both bacteria and virus versions of WIMP. Here we include any read mapping to *Escherichia coli* or a strain thereof as Ecoli, and any read mapping to *Lambdalikevirus* as Lambda, although the algorithm was not able to identify reads beyond the genus level for viruses.

ONT has introduced a graphical interface titled “What’s in my Pot”” (WIMP) for the Kraken taxonomic classifier, which uses kmer alignments to deter mine sample identities down to the strain level^11^. At the time of running, Metrichor™ offered separate pipelines for bacteria and virus classifications via WIMP, although a combined version is now available. The pipeline attempts to classify only 2D reads that pass a certain quality threshold. We ran bacterial and viral classifications separately on the ground data to compare to our BLAST results (Figure 3b). WIMP “Bacteria k24” identified 88 reads classified over its default threshold score of 0.005, 87 of which were classified as *Escherichia coli*, with 12 of those further identified as particular strains of that species (although none matched the correct K1 2 strain). The final read was classified as *Methanosarcina barkeri str. Fusaro*. WIMP “Viruses k24” classified 37 reads under the genus *Lambdalikevirus*, and was not able to provide any further details. In total, 124 reads were classified as *E. coli* or *Lambdalikevirus* by the two versions of WIMP, a number consistent with the 2D BLAST results. With low query coverage and identity using BLAST, and given WIMP is only currently applicable to 2D reads for separate subsets of species, we explored two alternative approaches inspired by music processing that directly use event space data.

### Uncovering Nanopore’s Fingerprints of Genomes (UNFOG)

Our first approach, titled UNFOG, attempts to construct fingerprints of reference genomes and reads based on their most informative frequencies over time. This employs methods from Shazam, a popular application (app) that identifies songs based on short clips of user input^12^. Shazam first fingerprints the reference collection by pairing peaks from the spectrogram of each song and storing the time between these peaks and the time offset from the beginning of the song. The algorithm then attempts to match similarly constructed fingerprints from the user input. Clips are classified based on the number of fingerprints that match a particular song at a consistent time offset.

We converted reference genomes into event space using the mean currents for each 5-mer stored in Metrichor^TM^’s hidden Markov model. As seen in Figure 2a, the amperages associated with various kmers in our real data followed a very similar distribution to the mean currents of the model. As in Shazam, we were able to construct fingerprints using peaks in a spectrogram of the signal (Figure 4).

**Figure 4:**
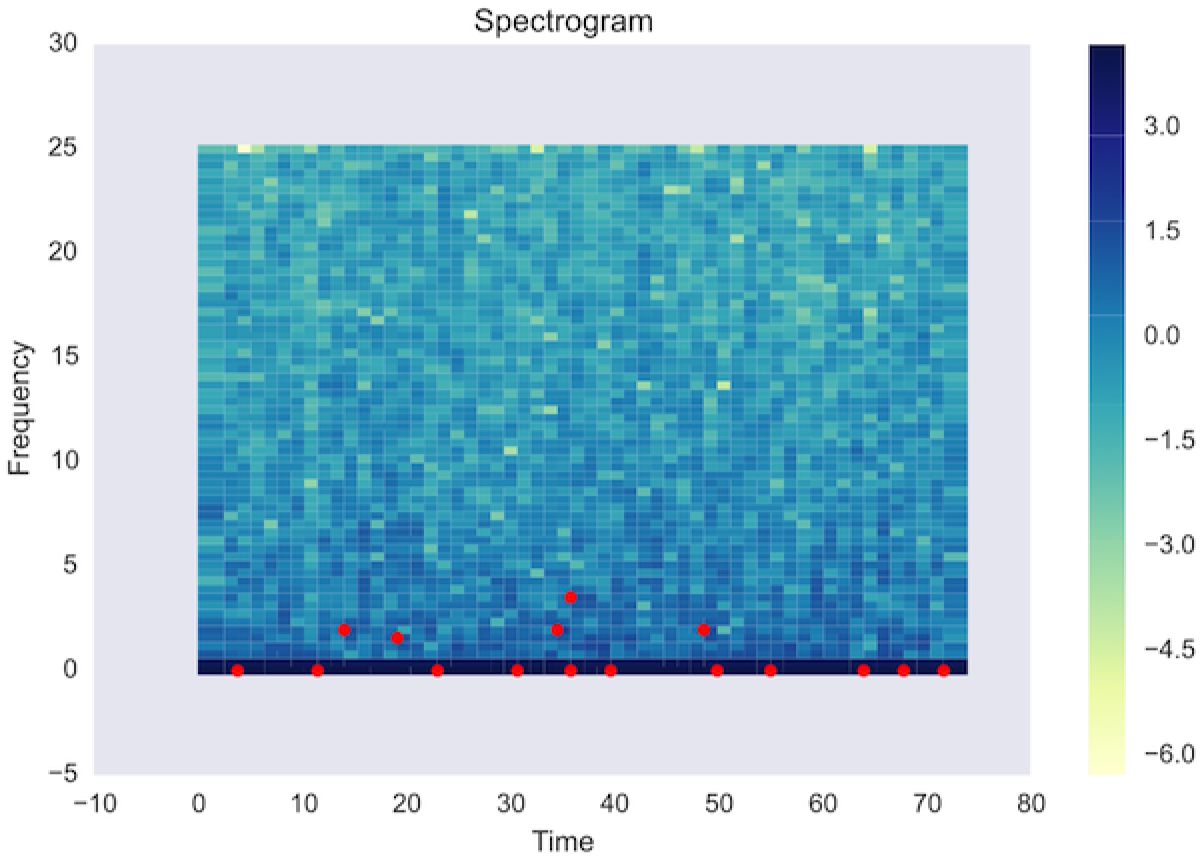
Spectrogram of the second flight data read (length = 1752) showing frequencies over time. Parameters used for the fast Fourier transform included a sampling rate of 50, a window size of 128, and an overlap between windows of 64. Peaks with amplitudes over a certain threshold (here, 1.3 on the log10 scale, marked in read) are identified and paired to construct fingerprints.

We first ran a series of benchmarking tests using a subset of 12 reference bacterial genomes, including highly related species, to determine how well the algorithm is able to classify fragments from a reference genome (*Enterobacter cloacae*, Figure 5a). The best version of UNFOG was able to correctly identify a sample read over 65% of the time and was largely tolerant to up to 10% mismatches. However, the percentage of reads correctly mapped dropped dramatically in response to induced insertions or deletions in the read. This is similar to what has been found in music identification, where fingerprinting algorithms fail to identify alternative versions of songs due to timing differences, and poses a particular issue for nanopore sequencing. The largest percentage of errors has been found to be deletions, at roughly double the rate of insertions or mismatches in 2D base-called sequences^13,^ review

**Figure 5:**
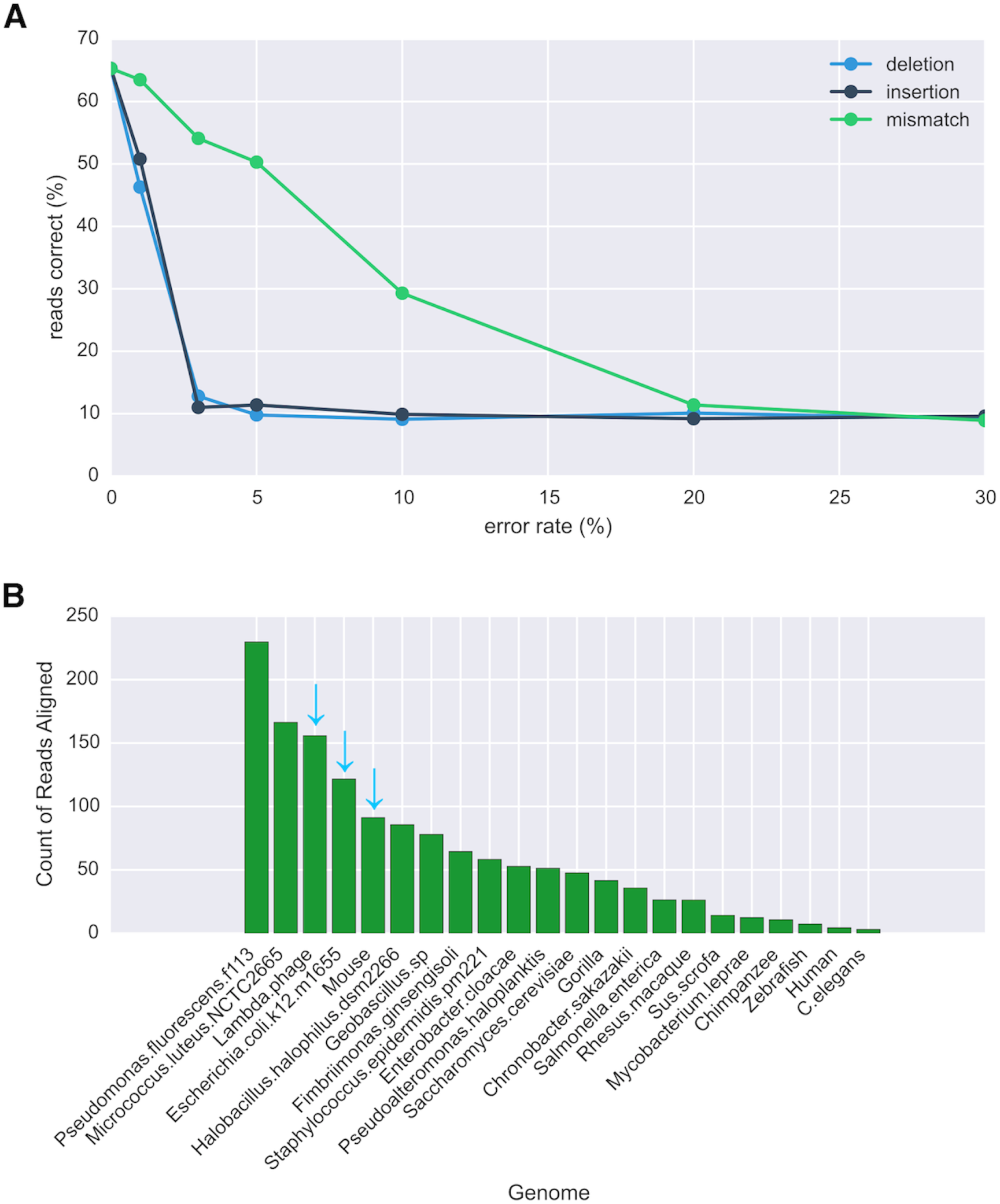
**a)** Benchmarking results for UNFOG using 5000 base pair fragments of reference genomes. The percent of reads correctly identified as the reference species was 65%, but dropped precipitously as we induced errors into the fragments, with greater tolerance to mismatches than to indels. **b)** UNFOG results for the ground data. *E. coli* was the top hit, as for BLAST and WIMP using 2D reads, and all three species in the sample were in the top 5 hits among 22 reference species.

Tests on the microgravity data failed to classify any of the reads as *Lambda phage, E. coli*, or mouse, identifying one read at *Staphylococcus epidermidis* and a second as either *Halobacillus halophilus* or human. Running UNFOG on ground data revealed more promising results, with all three species in the top 5 of 22 reference genomes, although *Pseudomonas fluorescens* and *Micrococcus luteus* ranked higher (Figure 5b).

### Uncovering Nanopore’s Signal Mapping over Genomes (UNSMOG)

Our second approach attempts to uncover greater similarity between sequences by converting them to entropy space, estimating the entropy for the signal using a generalized correlation integral approach^14^. This approach has also been effective in song identification, and has correctly identified the same songs across versions by different artists^15^. We had previously found that reads showed visually similar entropy patterns to the reference sequences of their BLAST hits. Initial tests with the flight and ground data revealed no such relationship (Figure 6a). However, we suspect that the high rate of stays in most of the reads from these data distorts the signal beyond the limits of this method. Once we reduced our search to reads with stay rates of <0.1 stays/base called, visual similarity was again apparent between the sequences in entropy space, including in areas where BLAST failed to align the sequences (Figure 6b). Sequences that fit the criteria of having a BLAST hit with relatively high query coverage and low stay rate were particularly scarce for this data set, but we had not previously observed this issue with other experiments.

**Figure 6:**
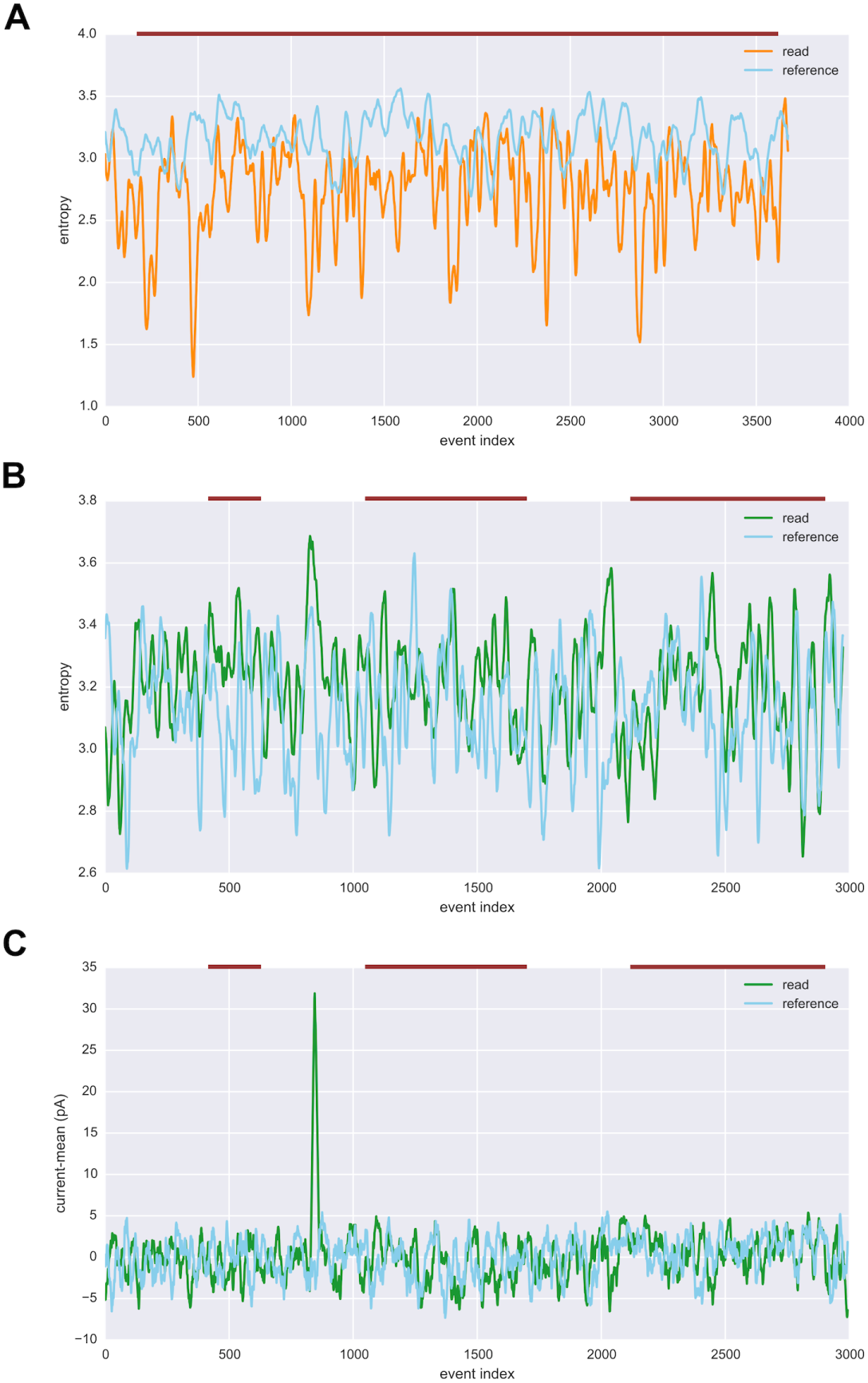
**a)** The flight data read with the highest mapping score and the subset of *Escherichia coli* genome to which it aligned in entropy space. The reference sequence is shown resampled to match the length of the read, with both signals smoothed by taking the mean over a sliding window of 20. Similarity was found to be 0.01 (p = 0.5) between the entropy signals using Pearson’s correlation. The approximate region of alignment for the BLAST hit is marked as the burgundy line above the plot, although indices do not correspond between event space and the base-called sequence due to the presence of stays and skips in the event space read. **b)** Entropy signals for a ground data read with a stays per base called ratio of 0.0009, far lower than the mean for these data (2.11). Query coverage for the BLAST alignment of this read to *Lambda phage* was 58%, with 68% identity. The cross correlation between the entropy signals after smoothing was low, r = 0.15, but positive and significant (p = 1.25×10^−17^). **c)** Similarly smoothed event signals for the same ground data read and reference sequence as shown in **b)**, which resulted in lower correlation, r = 0.04 (p = 0.06), suggesting that taking the signal entropy may help match event space representations to references.

## Discussion

Nanopore sequencing promises to contribute to healthcare in increasingly remote settings, as recently demonstrated during the Ebola outbreak in West Africa^16^. We show here the possible beginnings of a new frontier in genomics and genetics, in which humans and robotic rovers will be able to bring DNA sequencers as they travel beyond low Earth orbit. Despite some difficulties in adapting the library loading protocol for the MinION™ sequencer to a microgravity environment of a parabolic flight, we successfully produced the first ever DNA sequencing read in a reduced gravity environment.

We propose several potential modifications to the instrument to facilitate use in reduced gravity. We observed that the attached MinION™ cover was difficult to control in microgravity when attempting to load the sample. We recommend that it have a catch to keep it in the open position during loading. Aboard the International Space Station (ISS), this would most easily be achieved with Velcro. The most important thing to consider when loading a sample is that bubbles are extremely difficult to deal with in microgravity and easy to create. Currently, if the pipette tip is not angled perpendicularly to the loading pore, it becomes impossible to both remove the initial air bubble and introduce sample into the pore. The sample will pool on the outside of the flow cell rather than entering the pore; instead of removing the initial air bubble you will pipette ambient air. Modifications to the flow cell design permitting wider deviations to the angle of pipette tip entry would facilitate proper loading. The crew procedure for use aboard the ISS will emphasize a completely perpendicular loading and firm pressure to completely seal the pore with the pipette tip before loading with the current flow cell version. Any extra space will allow the liquid to pool on the flow cell surface.

We also observed leakage from a vent on return to the Johnson Space Center. In the microgravity conditions on the ISS, leaking due to rotation or tilting of the sequencer is extremely unlikely to occur, because surface tension and cohesion should be the dominant forces. Nevertheless, to prevent leakage in the future during the transport of flow cells after they have been loaded with samples on Earth, we recommend a gas-permeable barrier for vents. Alternatively, if venting only occurs during sample loading, a completely closed vent cover could be devised that is controlled by the sliding door covering the sample loading pore such that when the door is opened, the vents also open. Once the sample has been loaded and the door closed, the vents could close as well.

In terms of running the event-calling software MinKNOW™, there were no issues with communication between the MinION™ and the tablet during the flight. A screen protector for the tablet that minimizes glare is recommended, as well as increasing the brightness setting and magnification of text and icons as much as possible. An attached rubber-tipped stylus would also be helpful as the buttons (even with high magnification) are quite small and close together. These issues could also be addressed through software changes, as the MinKNOW™ software has not been optimized for use on the Surface Pro 3 or the Windows 10 operating system we employed. Based on our observations, the primary issue with the sequencing was a low number of available pores for sequencing. As mentioned previously, the flow cell used for in-flight sequencing was outside its optimal use period. A second potential factor reducing nanopore availability may have been caused by the parabolic flight itself, as the gravity conditions oscillate between microgavity (during freefall) and 1.8G (during ascent for the next freefall). These changes in gravity regimes could create more opportunities for air bubbles to migrate into the nanopores to obstruct them. We note that in this regard, the parabolic flight environment is different from the ISS, where the flow cells will experience 1G on Earth, a brief interval of enhanced gravity during launch, and then microgravity for until they are disposed of after use on the ISS. We hypothesize that the MinION™ would perform more comparably to a ground control setup when operated in a constant microgravity setting such as the ISS.

The reads produced during the flight demonstrated several distinguishing traits from those produced during the ground control experiment. Currents were consistently higher in the flight data, although it is difficult to ascribe this change a particular source or sources without changing some of the underlying circuits of the instruments and testing them again. We have previously observed deviations between different data sets, but they were often run on different versions of the chemistry, confounding the comparisons. Notably, noise was only significantly higher in one flight data read, although the number of stays across events was high in all three reads as well as in the ground data.

Our tests of WIMP, the online metagenomics classifier, demonstrated several potential issues. First, users cannot yet choose their reference databases beyond bacteria, viruses, or bacteria, viruses, and funghi. In addition, though classifications are stored in the fast5 files for each read, there is no way to save the final report from the GUI. This report would be helpful for quicker reference and comparison to other methods. WIMP also classifies only 2D reads, which could prove problematic for metagenomics studies where the number of reads for an individual species may be quite low to begin with, and where the 1D reads would be useful as well.

Our work has also shown that new methods may be needed for processing of these data. Both WIMP and BLAST rely on successful base calling by Metrichor™. With particularly noisy data, BLAST can fail to align a majority of reads. Though for our data BLAST was usually able to identify several candidate species, their scores are not necessarily high or distinct enough to permit positive identification.

While we show that UNFOG has potential in theory, the nature of errors in nanopore sequencing currently limit its application. An entropy-based solution may be capable of greater accuracy at a significant cost in computational time; however, as we demonstrate here, that this may fail for reads that are extremely stretched out in time with respect to their reference sequences. Several programs attempt to deal with an analogous problem in music, that of “query by humming”, where, far from the exact versions of songs Shazam requires, a user can identify a song by humming a short segment of melody^17,18,19^. As the chemistry continues to improve and error rates decrease, we suggest that adapted methods for fingerprinting could allow for rapid metagenomic classifications using future iterations of nanopore sequencing technology. The greatest advantage to fingerprinting would be speed: UNFOG was able to classify the almost 2000 reads from the ground data in less than three minutes after the database was built.

## Methods

### Sample preparation

Each of the three types of genomic DNA samples was prepared for sequencing according to the procedures specified by ONT, beginning with 1 *μ*g of each sample of organismal DNA (bacteriophage lambda, *E. coli* and mouse). To facilitate sample loading during the microgravity intervals of the flight, we deviated from the recommended three-step sample-loading procedure, which entails loading 150 *μ*l of running buffer containing fuel mix, waiting 10 minutes, repeating the previous two steps, and completing the procedure by loading a final 150 *μ*l of the complete sequencing mix. Instead, we preloaded syringes each containing the full 450 *μ*l volume of running buffer and fuel mix supplemented with 6 *μ*l of pre-sequencing mix from each organism. The libraries and pre-loaded syringes were prepared two days prior to flight, stored at −20°C until the day of the flight, and stowed at ambient temperature aboard the plane prior to loading the flow cell.

### BLAST

Nucleotide-Nucleotide BLAST 2.2.29+ was run using default settings for somewhat similar sequences, connecting to NCBI’s most recent nt database. Blast hits were preferentially counted towards the sample species even if others scored higher. If there were hits for multiple sample species only the highest scoring was considered. Query cover was calculated for the counted hit by taking dividing the length of the primary alignment by that of the query sequence. This differed at times from the query coverage calculated using the online version of BLAST, which is able to calculate over an aggregation of compatible aligning regions, and thus found a query coverage of 26% for the longest flight data read in its alignment to mouse. Identity was as provided by the BLAST alignment report.

### WIMP

Base-calling and species classification were performed using the WIMP Bacteria k24 for SQK-MAP005 (version 1.27) and WIMP Viruses k24 for SQK-MAP005 (version 1.27) pipelines. Species counts were combined for Figure 3b, with reads mapping to the genus *Lambdalikevirus* considered positive hits for *Lambda phage* in the absence of species-level classification for the Viruses version. We considered hits only above the default threshold score in the GUI; decreasing that threshold to zero showed more false hits for the Bacteria run and had no effect for the Viruses run.

### UNFOG

After testing multiple parameter sets, we chose a window size of 128 and overlap of 64 to compute the FFT. We tested several sampling rates; shown are the results using a sampling rate of 10 for the test data and 50 for the real data. Spectrogram peaks over an amplitude of 5 were paired if within 100 of one another on the time axis of the spectrogram for the benchmarking test version, this was changed for the real data version to an amplitude of 20 and time limit of 300 in an attempt to increase specificity. Paired peaks were stored as hashes, along with their offset times for retrieval and comparison.

The benchmarking tests were run using a thousand random 2500-base fragments of *Enterobacter cloacae* genome joined to their reverse complements to create 2D reads. Errors were induced prior to conversion of the reads from kmers to currents to mimic possible rates in output base-called sequences. The best version of UNFOG during testing involved storing all instances of a fingerprint across each genome and later removing from consideration any fingerprints that appeared over 50 times in any reference genome. However, this modification did not improve results for the real data (possibly because with high error rates, exact matches are unlikely to begin with) and significantly increased classification time, therefore a previous version that saved only the final instance of each fingerprint was used for the real data. In the future, a step discarding the most common fingerprints will likely be incorporated into building the database. For each read, the species with the highest number of matches at a consistent offset time from the beginning of the reference sequence was counted. With both template and complement strands, the same offset time had to be found for both strands for a positive hit. In the case of ambiguous matches, the count for each potential species was increased by 1/(number of matches).

Code for UNFOG is posted at http://pbtech-vc.med.cornell.edu/git/mason-lab/unfog.

### UNSMOG

Entropy was calculated for each base using a generalized correlation integral approach with a sliding window of 20 and overlap of 19^14^. We then smoothed the signal by taking the mean over a similarly sized sliding window.

